# Application of Coincidence Index in the Discovery of Co-Expressed Metabolic Pathways

**DOI:** 10.1101/2023.06.26.546540

**Authors:** João Paulo Cassucci dos Santos, Odemir Martinez Bruno

**Affiliations:** Scientific Computing Group, São Carlos Institute of Physics, University of São Paulo (USP)

**Keywords:** Systems Biology, Halobacterium salinarum, Coincidence Index, Graph Theory

## Abstract

Analyzing transcription data requires intensive statistical analysis in order to obtain useful biological information and knowledge. A significant portion of this data is affected by random noise or even noise intrinsic to the modeling of the experiment. Without a robust treatment, the data might not be very thoroughly explored and even incorrect conclusions could be drawn. Examining the correlation between gene expression profiles is one of the ways bioinformaticians extract information from transcriptomic experiments. However, the correlation measurements traditionally used have worrisome shortcomings that need to be addressed. This paper compares the two most common correlation measurements, Pearson’s r and Spearman’s r, to the newly developed coincidence index, a similarity measurement that combines Jaccard and interiority indexes and generalizes them to be applied to vectors containing real values. We used experimental data from a microarray experiment from the archaeon *Halobacterium salinarum* that evaluates the effects on the organism when exposed to light in an anaerobic environment. The utilized method explores the co-expressed metabolic pathways by measuring the correlations between enzymes that share metabolites and searches for local maxima using a simulated annealing algorithm. We demonstrate that the coincidence index extracts larger, more comprehensive, and more statistically significant pathways than the traditional Pearson’s and Spearman’s measurements.

## 1 Introduction

The use of transcriptomic analysis to better understand the metabolism of living organisms has become standard practice in molecular biology as the cost of these experiments significantly decreased Liang (2013). This has generated a large amount of biological data, necessitating improved methods for analysis and information extraction. One way to perform this task is to elucidate biologically correlated genes, using a guilt-by-association approach where similar expression profiles suggest a shared molecular or biological function Wolfe et al. (2005).

In conjunction with this method, enzyme-enzyme interaction graphs based on known shared metabolites are used as an additional biological evidence of correlation.Ideker et al. (2002); Patil and Nielsen (2005) Such graphs have their edges weighted according to the correlation measurements of the expression profiles. The use of known enzymatic reactions can reduce the number of gene profiles to be correlated, can provide stronger evidence for common biological function, and eliminate the need for all genes to be enriched. Instead, only the enzymatic sub-network, which is much more relevant for understanding the organism’s metabolism, needs to be enriched. Patil and Nielsen (2005)

Pearson’s correlation coefficient is the most common metric used to evaluate the similarity between two vectors. However, it can be ineffective against noisy data due to the presence of outliers that can change the result of the measurement significantly. Spearman’s correlation coefficient attempts to mitigate the effects of these outliers by ranking the elements of the vectors, but this approach has limited effectiveness.Patil and Nielsen (2005); Schober et al. (2018)

The coincidence index is a new measure of similarity between vectors defined by the combination of the Jaccard Index and Interiority Index.da Fontoura Costa (2022) Traditionally, these indices are used to compare sets containing discrete elements. However, by employing the concept of multi-sets, they can be generalized for application to vectors containing real values.Blizard (1991)

This work aims to provide a comparative analysis between these three measurements. The objective is to show the differences in the application of each and in the biological conclusions that can be drawn from them. The biological experiment used for this comparison is a microarray transcriptome time series of the Archeae *Halobacterium salinarum*, a photosynthetic and halophilic organism that has a very well-characterized metabolism due to its dynamic energy extraction pathways.Gonzalez et al. (2008) The experiment in question exposed the microbe to an anaerobic condition with continuous light exposure for 72 hours and collected samples every 3 hours.Baliga et al. (2004)

## 2 Materials and Methods

### 2.1 Transcription Data

The transcription data comes from a microarray experiment in which *H. salinarum NRC-1* was exposed to continuous light in an anaerobic environment for 72 hours. The experiment, obtained from NCBI, has the series GEO accession number GSE7712. It contains four spots for 2400 unique gene sequences and employs a dye-swap technique to correct for dye bias.Bonneau et al. (2007) Samples were collected every three hours, resulting in 25 time-points in the series. To obtain more reliable results, we calculated the median value of the 4 samples for each of the 468 genes contained in the network to avoid the effect of outliers. Genes present in the experiment that were not part of the network were disregarded. The data points in the experiments included a flag indicating the reliability of each spot’s measurement: ”I” for reliable measurements, ”J” for intermediate ones, and ”K” for unreliable ones. Spots marked with a ”K” flag were discarded.

### 2.2 Building the Enzyme Interaction Graph

The enzymatic reactions to construct the Enzyme Interaction Graph were obtained from a characterization of the *Halobacterium salinarum* genome. This reconstruction includes a total of 559 reactions involving 468 ORFs, all of which code for enzyme synthesis (Gonzalez et al., 2008). The approach used in building the network is the same as that described in Patil and Nielsen (2005) and is illustrated in Figure 1. The nodes of the graph represent the 468 enzymes, and the edges between them are established based on whether they share a metabolite. If they share at least one metabolite, an edge is established between them.

**Figure 1.**
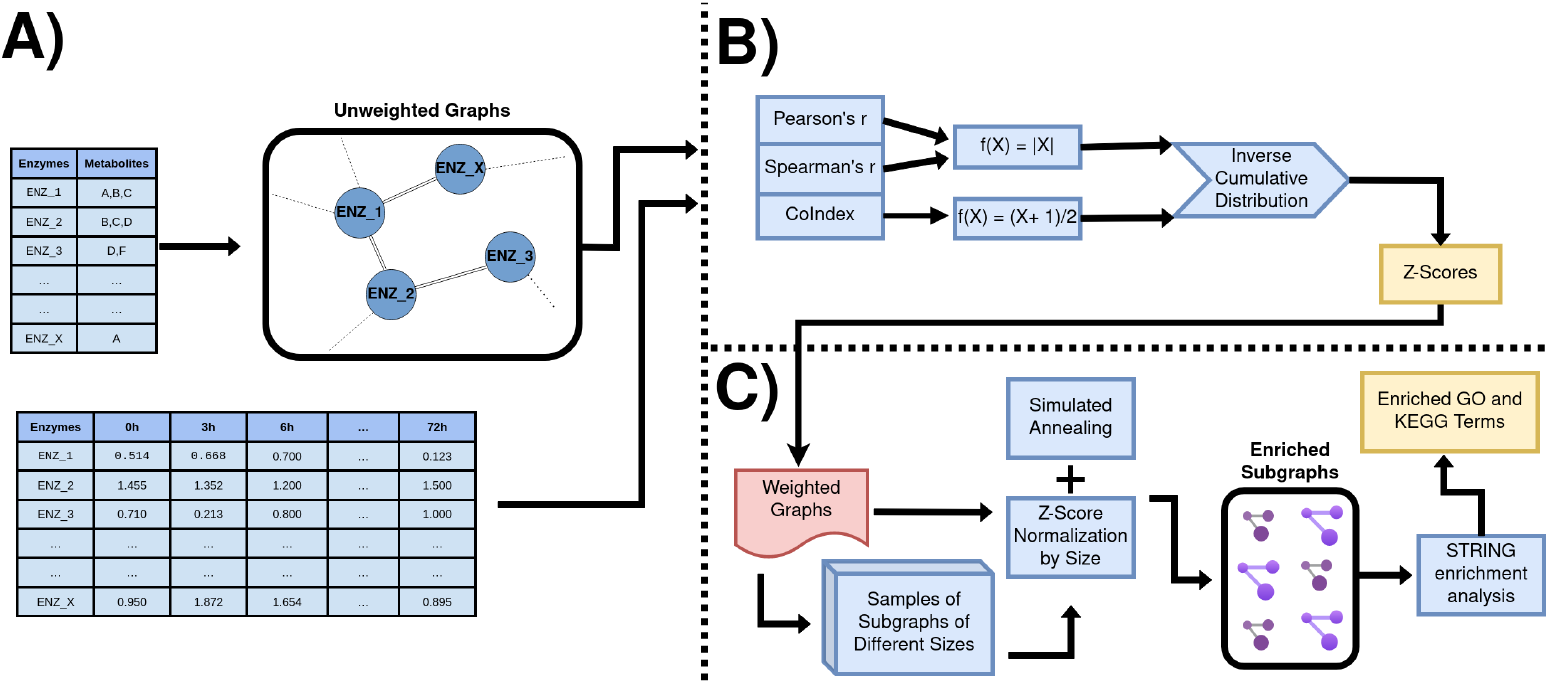
General pipeline of the methods used in this work. A)Construction of the enzyme-enzyme interaction graph, using the shared metabolites between them as the basis for the edges. The dataset containing the median values of the microarray experiment for each gene in a time-series is also shown. B) Depiction of the measurements used to obtain the weights of the edges as Z values, note the difference in the processing of Pearson/Spearman and coindex measurements. C) Normalization of the Z values based on their size, together with the simulated annealing algorithm in order to extract the local maxima (represented in the Enriched Subgraphs). These graphs then have their nodes searched and enriched in the STRING database in order to find relevant biological information associated with them.

The resulting network is highly interconnected, with the average shortest path between all nodes being less than 2. There are a total of 41266 edges and an average node degree of 176.35. Most of these edges are established from the ATP/ADP pair, the redox cofactors NAD+/NADH and the H2O/H+ pair. These molecules are fundamental to many metabolic reactions, they are referred to in molecular biology as ”energy tokens”. They can drastically influence the outcome of co-expressed enzyme extraction by connecting biologically unrelated enzymes through these common metabolites. For this reason, it is important to experiment with which metabolites to work with in order to obtain the most coherent results.

### 2.3 Usual Measurements

The measurement used to compare two expression vectors in the method described by Patil and Nielsen (2005) is the Pearson correlation coefficient, also known as Pearson’s r. This is standard practice, as Pearson’s r is the most commonly used correlation coefficient. However, it has limitations. It can be very susceptible to overestimation in noisy data due to its sensitivity to outliers, and microarray data tends to be very noisy.Schober et al. (2018) The method calculates the Pearson’s r value of the expression vectors of enzymes/nodes that share a metabolite/edge.

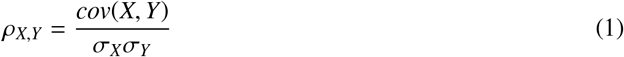

Pearson’s r produces values in the range from -1 to 1, with -1 being a perfectly linear inverse correlation and 1 a perfectly linear direct correlation. To evaluate the significance of these correlations, an inverse cumulative distribution is applied to the absolute value of Pearson’s r to obtain Z values for the edges (*Z*_*es*_) in a normal distribution. These Z values, which range from negative infinity to positive infinity, are used to determine how many standard deviations a particular measure is from the mean.

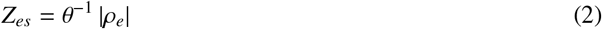

Another traditional correlation measurement, albeit more computationally intensive is Spearman’s r. This measurement ranks the instances within the vectors 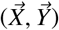 and then performs the same calculation as Pearson’s r. This approach circumvents the requirement of Pearson’s r for a linear correlation between instances. Instead, Spearman’s r quantifies the correlation between any monotonic increase or decrease in the vectors.

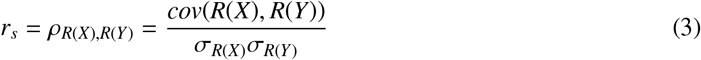

### 2.4 Coincidence Index

The coincidence index (coindex) is a recently developed similarity coefficient that has shown promising results in complex network analysis. It is the product of the Jaccard and interiority indexes (the interiority index is also known as the overlap coefficient).da Fontoura Costa (2022) Traditionally, these indexes are employed using the properties of sets, whose elements are all distinct and comparable only to their equivalents. However, by employing the multisets concept, it is possible to generalize the Jaccard and interiority indices to real-valued vectors.da Fontoura Costa (2022)

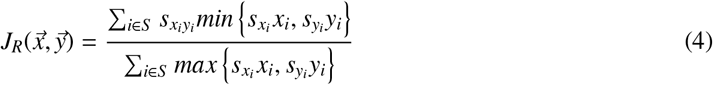

In equations (4) and (5), *x*_*i*_ and *y*_*i*_ correspond to the i-component of the vectors 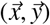 of size S, 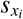 and 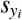 correspond to the sign of *x*_*i*_ and *y*_*i*_ and 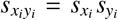. These real-valued indexes are then multiplied together to form the coindex.

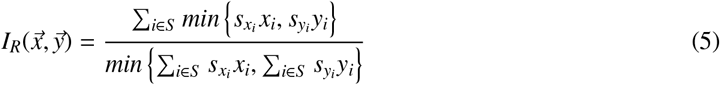

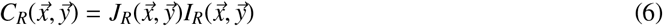

Coindex has been shown to provide a more strict and detailed measurement of similarity between vectors compared to more traditional measurements.da Fontoura Costa (2022) However, it is important to note that the interpretation of its output differs from that of Pearson’s and Spearman’s r. The output values also range from -1 to 1, but the minimum value represents total dissimilarity between the vectors, rather than a perfect inverse correlation. This makes equation 2 inappropriate for this measurement, as it could conflate vectors deemed very dissimilar with those highly similar to each other. It could also skew the distribution of the values towards 0, generating an asymmetrical Gaussian distribution that distorts the meaning of the Z values. For these reasons, we chose to map the coindex outputs in the following way:

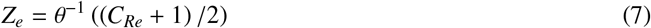

In Figure 2 the coindex is shown to be more stringent in every scenario involving random variables than Pearson’s and Spearman’s r. It not only is more sensitive to noise than the others, but it also shows a greater sensitivity to non-linearity (more so than Pearson’s r) and to variations in scale, aspects that the other measures do not account for.

**Figure 2.**
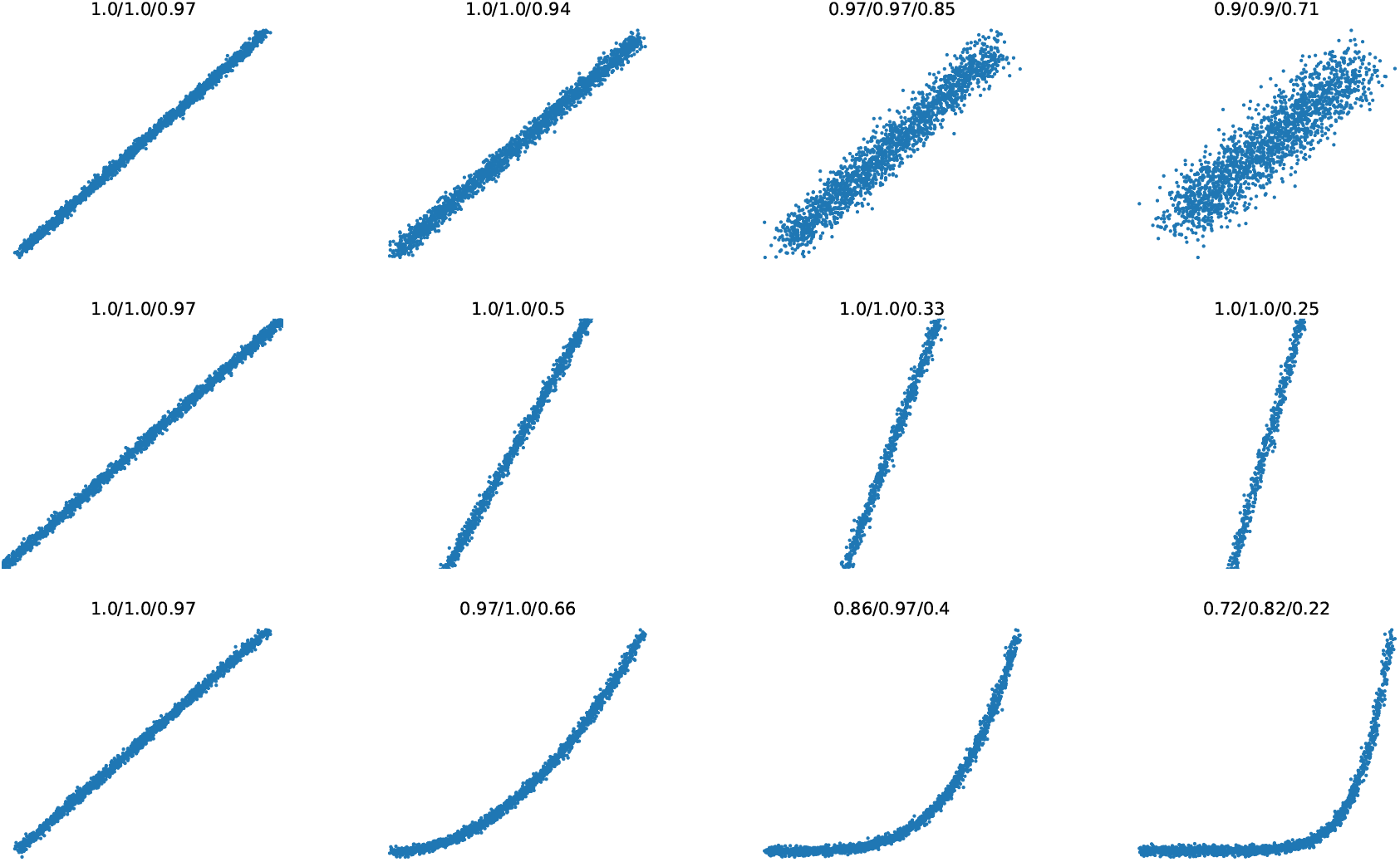
Comparison of the three measurements using two example vectors, X and Y, with 2000 values each. The values on top of each curve represent Pearson’s r, Spearman’s r, and coincidence index respectively. The first row shows a linear correlation between X and Y with more random noise in each iteration. The second row shows a very strong linear correlation but with a larger angular coefficient in each iteration. The third and last row shows increasingly non-linear correlations between the vectors.

### 2.5 Scoring the subnetworks

Our goal was to find highly correlated/similar subnetworks that can provide insights into the mechanisms being co-expressed by the archaeon. To achieve this, we performed the same calculation described in Patil and Nielsen (2005) to score a subnetwork of size k:

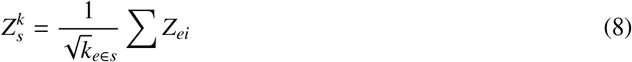

Here, *Z*_*ei*_ represents the Z value of each edge in the connected subnetwork. However, this value needs to be adjusted according to the background distribution of the 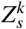 value from other connected subnetworks of the same size k. This adjustment is necessary to ensure that larger subnetworks do not end up with larger Z values due to their greater prevalence. This adjustment can be made as follows:

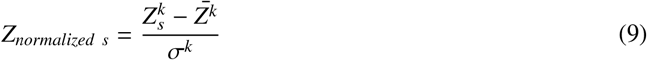

With this measurement, it is possible to determine whether a resulting subnetwork is affected by the experimental conditions more than expected without a bias towards size. The average 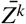 and the standard deviation σ^*k*^ can be obtained by sampling connected subnetworks within the main network. In this way, we can obtain a certain amount of samples (around 200) with which we calculate the mean and standard deviation of the Z values of size k subnetworks to normalize for size variations. Assuming a network has *N*_*max*_ nodes in total in its largest connected component, there are *N*_*max*_ − 1 different possible connected subnetwork sizes, and thus, *N*_*max*_ − 1 means and standard deviations. Subnetworks of size k=1 don’t have edges, therefore they cannot be scored.Baliga et al. (2004)

The flowchart presented in Figure 3 exemplifies how the mean and standard deviation were obtained through a sampling of random connected components of size k.

**Figure 3.**
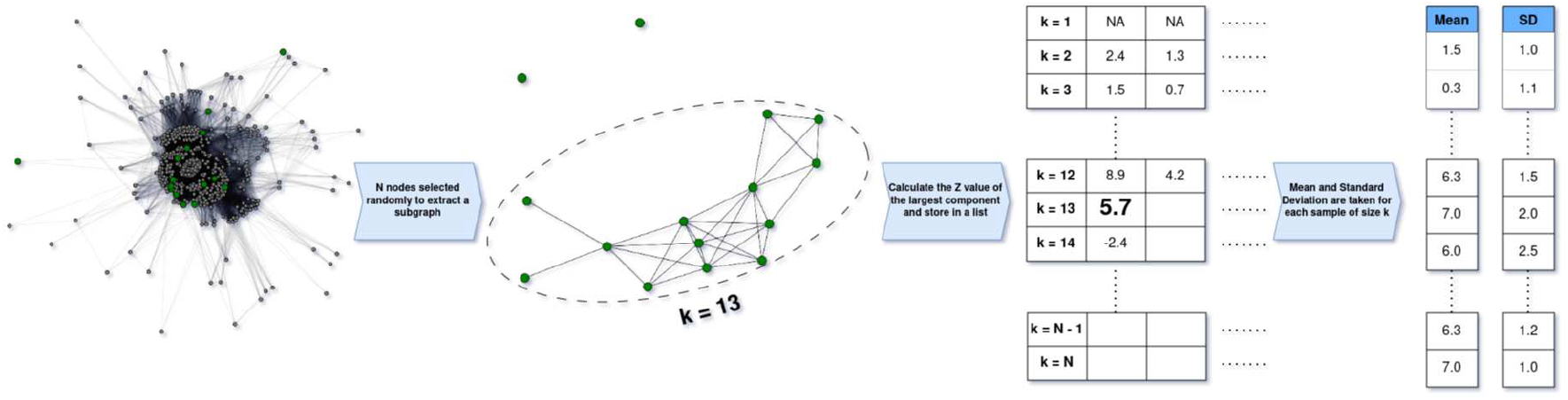
Sampling method used to perform the normalization of the connected network Z values according to their size. First, N nodes are selected randomly, after that, the largest connected component extracted from these nodes has its Z value calculated. The Z value is then stored in a list together with other samples of the same size k. With the samples collected, the mean and standard deviation of every list is calculated, generating the two lists that will be used in the standardization of the networks. This is done 200 times for each possible value of N ranging from 2 to *N*_*max*_

To identify enriched subnetworks within the main enzyme-enzyme interaction graph, we used the simulated annealing algorithm Patil and Nielsen (2005); Ideker et al. (2002). This method performs a heuristic search for local maxima within the main network, finding groups of connected genes with a higher-than-average *Z*_*normalized s*_. The method initially assigns a state of 1 or 0 to the nodes of the network at random, then ‘masks’ the nodes in the 0 state and searches for the highest-scoring connected component (or connected subnetwork) among the nodes in the 1 state. Once the highest-scoring component is found, the algorithm randomly toggles a node’s state and performs the search again. If the score is higher than before, the toggled state is maintained; if it is lower, the probability of maintaining the toggled state is determined by the function 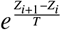, where T is the current ”temperature” of the system. This function outputs a value between 0 and 1.

If the value of the output is smaller than a random number from [0,1], then the node is untoggled; otherwise, it remains toggled. This process helps to avoid ”false maxima”, allowing the algorithm to perform a search more thoroughly on the network.

This search is performed N times, with the value of the temperature T decreasing geometrically from *T*_*i*_ until *T* _*f*_ = 0.01. After this, a new search is conducted with T = 0 to ensure that the local maximum is found (a process known as simulated quenching).

We used some heuristics described in Ideker et al. (2002) to obtain a better result. One of these heuristics involves searching for a number M of connected components simultaneously, maximizing the sum of the score of these components instead of their individual Z values. This approach has generally been shown to yield better results. Ideker et al. (2002); Patil and Nielsen (2005)

Another heuristic involves an exception for nodes considered ”hubs”. These nodes can end up connecting many components together that are significant when separated but have a lower score when connected due to insignificant relationships that the hub introduces. To avoid these pitfalls, if a node exceeds a certain minimal degree value (*d*_*min*_), only the neighbors belonging to the highest scoring component that the hub is connected to are kept ”on” every time the hub node is turned ”on”. The neighbors that do not satisfy this criterion are set to ”off”. This minimizes the effect that hubs have on the simulated annealing process by restricting it to the component that has the largest score, thus avoiding penalization of smaller components.

Optimization of the parameters *T*_*i*_, *N, M* and *d*_*min*_ was carried out in order to get the best possible result by performing different simulations with varied values for them. Moreover, since the method is heuristic, it does not guarantee the optimal result, so 10 processes were run in parallel after choosing the best parameters. The process that yielded the largest Z value had its high-scoring components selected and analyzed.

### 2.6 Gene Ontology Enrichment

To determine whether the genes that compose each connected component contain any known biological information, a gene ontology enrichment analysis was conducted through the STRING database for all of the high-scoring connected subnetworks.Szklarczyk et al. (2022) The enrichment was done using only the genes from the larger enzyme-enzyme network as the background of the analysis. This approach helps us avoid drawing invalid conclusions based on incorrect a priori knowledge.Ashburner et al. (2000); and Seth Carbon et al. (2020)

The STRINGdb includes 445 out of the 468 genes in the main network. These genes were matched to the database’s corresponding identifiers and a GO term enrichment analysis with FDR enabled was performed for all of the components identified through the simulated annealing algorithm. The KEGG database was also utilized for the enrichment of biological pathways.Kanehisa et al. (2022)

## 3 Results and Discussion

### 3.1 Optimizing the parameters of the Algorithm

To obtain the best possible result, the simulated annealing algorithm needs to be optimized, as it is heuristic, non-deterministic, and multivariable. Furthermore, since we are experimenting with new similarity measures, it is important to determine which parameters better suit each one of them. To do this, we established ”default” parameters for the algorithm (N = 10^5^, m = 20, *T*_*i*_ = 1, *d*_*min*_ = 100) and varied one at a time. The parameter that yielded the highest Z value was then used to search the enriched subnetworks. The results can be seen in Table 1.

**Table 1:**
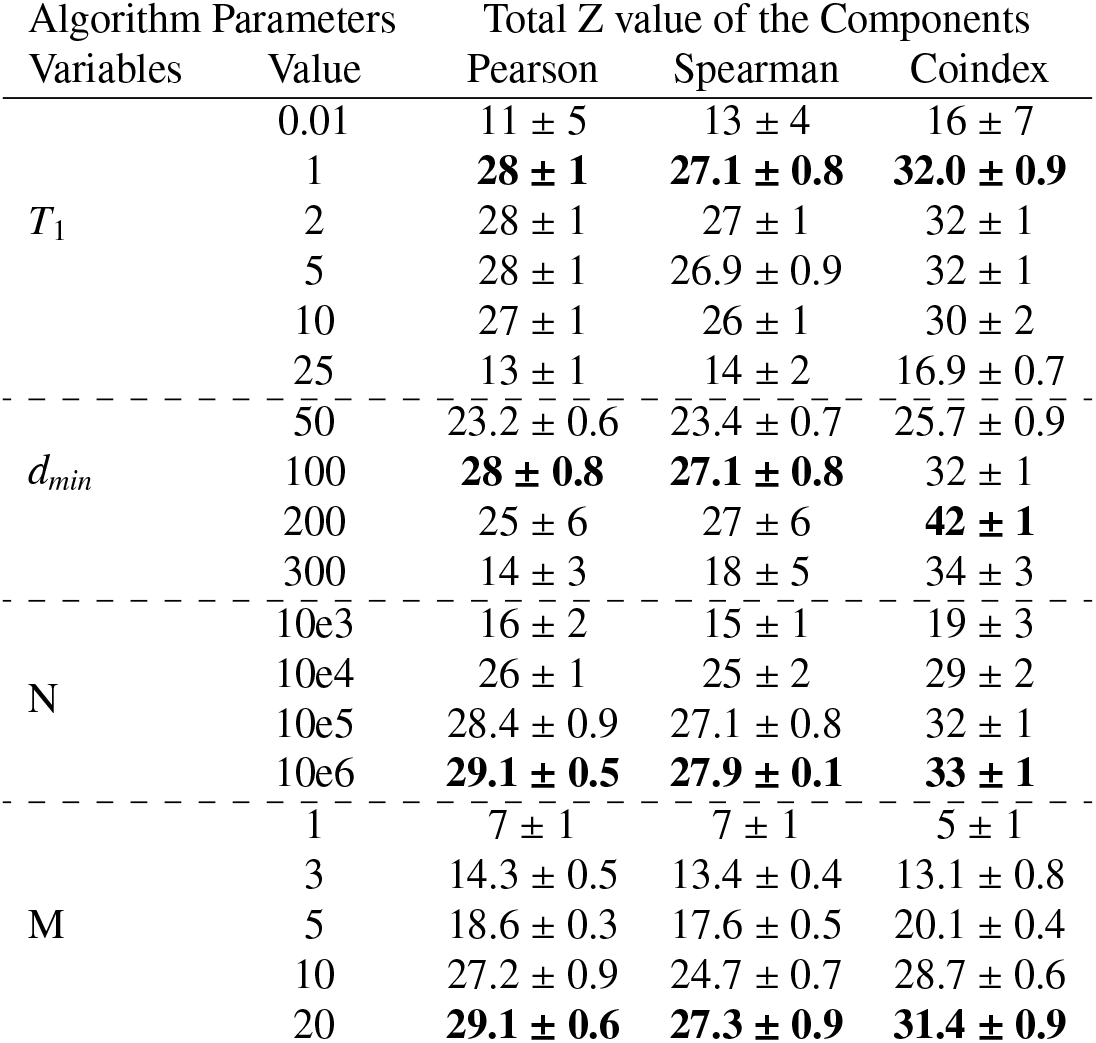
Values of the summation of the Z values from the connected components found with the respective changed value of the Variables. The highest values obtained are highlighted in bold numbers.

The table indicates that the optimal parameters for Pearson’s and Spearman’s r are the same (*T*_1_ = 1, *d*_*min*_ = 100, *N* = 10^6^, *m* = 20) and the resulting Z values are not significantly different either. The coindex, however, generally yields higher resulting Z values, and the optimal minimal degree value to consider nodes as hubs is *d*_*min*_ = 200. The variation in the optimal *d*_*min*_ implies that enriched coindex subnetworks are more connected and have more nodes compared to Pearson’s and Spearman’s r subnetworks.

The search for local maxima was also conducted in a second enzyme-enzyme interaction graph that excludes the most common metabolites. It has been demonstrated that some metabolites can disrupt the search for pathways by connecting biologically unrelated genes together, thereby establishing an excess of trivial connectionsGerlee et al. (2009). This also leads to the creation of many more hub nodes, necessitating a more careful search and, consequently, a more computationally intensive one.

To determine which ones to remove, we counted how many enzymes have reactions containing a particular metabolite. If the number of enzymes containing it exceeded 100 (roughly 1/5 of all the enzymes in the data), it was considered common and was removed from the network-building process (shown in Figure 4). There are more sophisticated methods that attempt to circumvent this procedure because it can be very arbitrary like chemical group tracking Huang et al. (2017). However, we are primarily interested in evaluating the performances of different similarity measurements in a network with a different topology, so this approach is sufficient.

**Figure 4:**
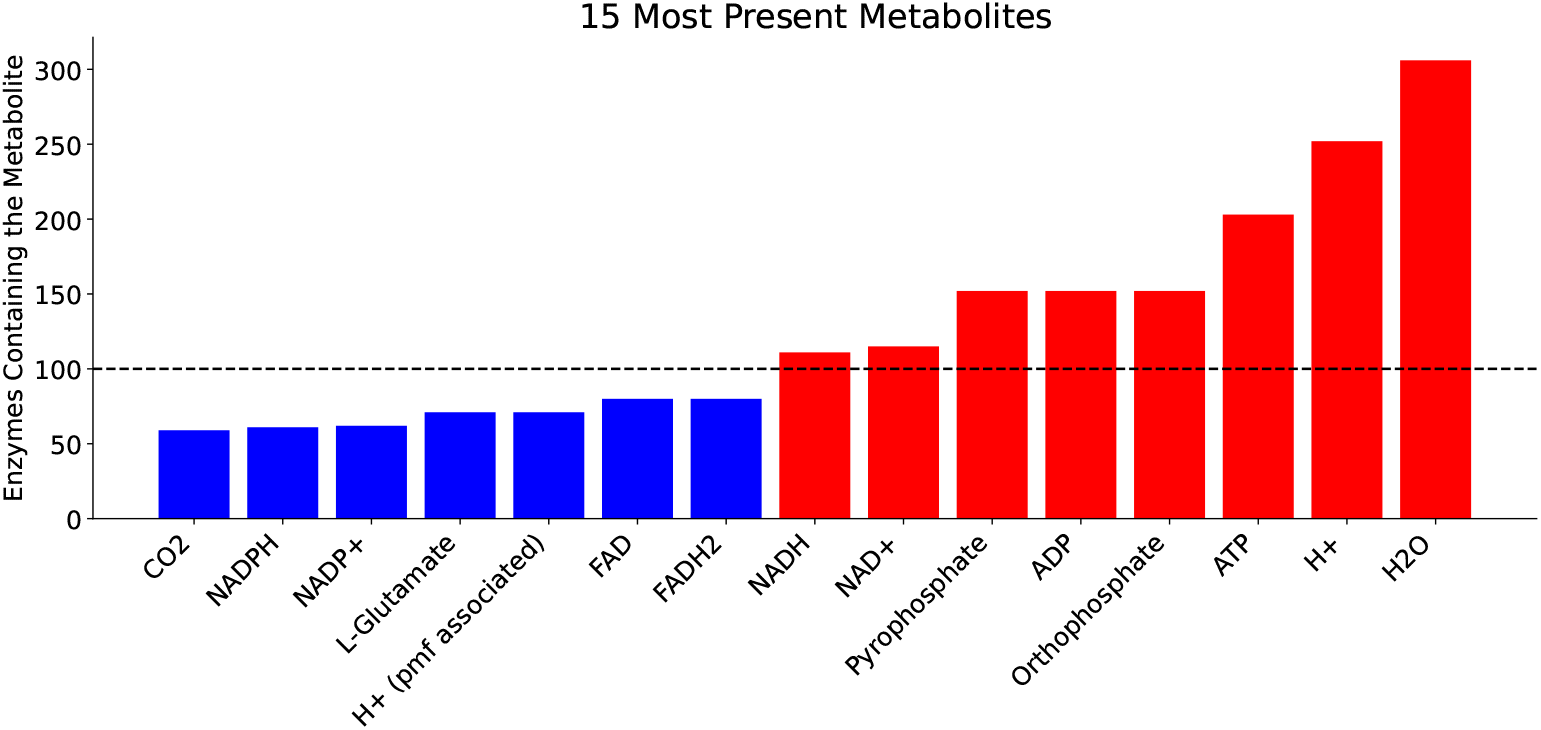
Top 15 most present metabolites in enzymatic reactions of the *H salinarum*. The ones present in more than 100 of the enzymes, highlighted in red, were ignored in order to build a second, less connected, graph.

As expected, the most common metabolites are primarily ”energy tokens” such as ATP/ADP. Actors in phosphorylation and hydrogenation were also among them. With their removal, the new graph becomes much less connected, with the total number of edges falling to less than a quarter of the original size and some nodes being removed due to the fact that they do not have any relationships to other nodes (shown in Figure 5). The average path length increased (from 1.69 to 2.68), but the new graph is still highly connected.

**Figure 5:**
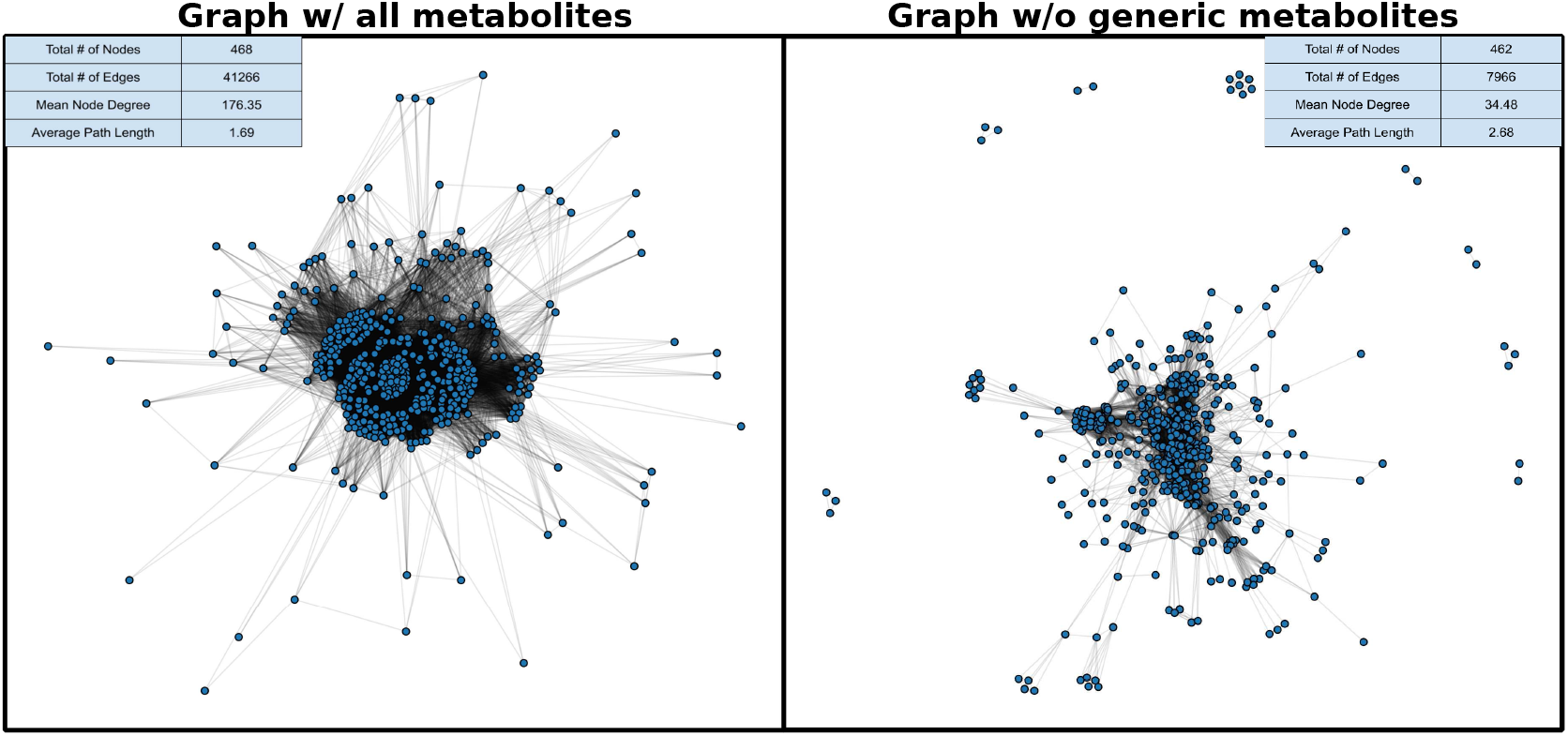
The two graphs used to obtain the enriched pathways. The one to the right contains all the metabolites involved in the Reactome of the *H. salinarum*. The one to the left is the graph with the most common metabolites removed. Some nodes are excluded because they don’t form edges with any other node, making them irrelevant. The number of edges decreases dramatically and the Average Path Length is bigger also.

Since the network’s topology has changed, its properties, such as the number of hubs and total connected components have also changed. As a result, we need to reoptimize the algorithm. This time, however, we only need to tune the parameters affected by changes in the number of edges, as the change in the number of nodes was not significant. These parameters are the number of connected components being searched simultaneously (M) and the minimal degree value to consider a node a hub (*d*_*min*_). The resulting Z values are shown in Table 2.

**Table 2:** Values of the summation of the Z values from the connected components found with the respective changed value of the variables. The highest values obtained are highlighted in bold.

| Algorithm Parameters |  | Total Z value of the Components |  |  |
| --- | --- | --- | --- | --- |
| Variables | Value | Pearson | Spearman | Coindex |
| dmin | 5 | 33.2 ± 0.3 | 34.1 ± 0.2 | 38.2 ± 0.3 |
| | 10 | 40.8 ± 0.9 | 42.8 ± 0.8 | 46 ± 1 |
| | 25 | **52.8 ± 0.8** | **51.6 ± 0.4** | 72 ± 1 |
| | 100 | 43 ± 4 | 44 ± 4 | **76 ± 5** |
| ----- |  | ----- | ----- | ----- |
| M | 5 | 22 ± 2 | 22 ± 1 | 42 ± 5 |
| | 10 | 36 ± 2 | 35 ± 2 | 58 ± 6 |
| | 20 | 54.0 ± 0.4 | 54.0 ± 0.7 | 79 ± 4 |
| | 40 | 68.9 ± 0.7 | 70.2 ± 0.2 | **91 ± 3** |
| | 80 | **69.0 ± 0.4** | **70.7 ± 0.7** | 91 ± 3 |

The optimal value for the minimal hub degree has decreased in all three measurements. This is expected since the mean node degree was reduced (Figure 5) resulting in hubs with a lower degree value. The coindex measurement, however, still maintains its larger optimal value, further indicating that the local maxima found by it tend to be more well-connected and contain more genes.

The optimal number of connected components to be searched simultaneously also increased in all three measurements, indicating that with the removal of the common metabolites, more local maxima have become evident.

### 3.2 Enriched Sub-pathways

After the parameter optimization, the algorithm was run again for all three measurements, this time, choosing the process that obtained the highest possible Z value. For the network containing all metabolites, all of the three measurements yielded around 10 enriched subnetworks but not all of them contained relevant information or had a significant Z value. The significance of the individual subnetwork was calculated through a one-tailed p-value test and a False Discovery Rate (FDR) correction. It is important to note that, since the normalization of the scores is done through a random sampling of all possible networks of a certain size, the value can vary in other simulations. However, for around 200 samples the variation of the Z values of the significant networks was below 5%.

Figure 6 illustrates how many components each measurement was able to find, and which ones include a group of genes that have biological information associated with them.

**Figure 6:**
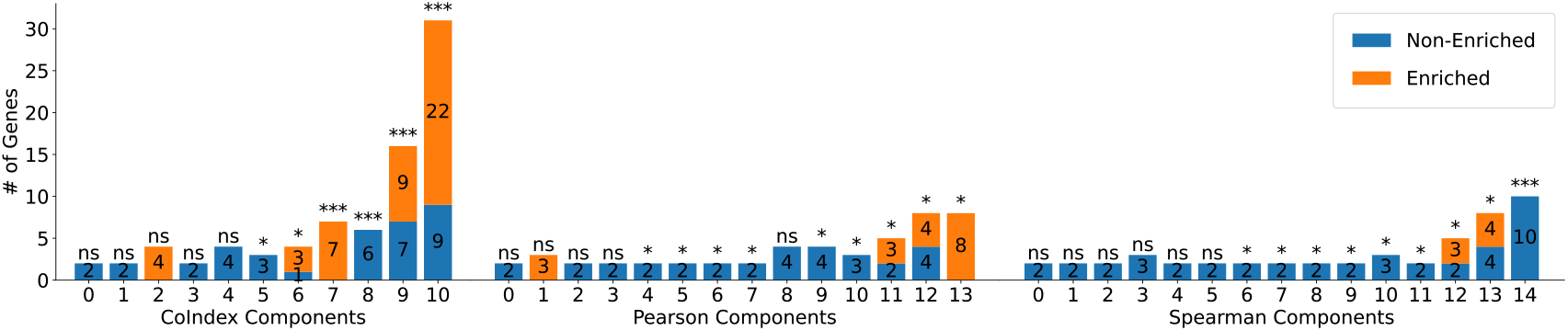
Barplots representing the number of genes contained in each connected component found by the Simulated Annealing Algorithm with tuned parameters. The genes that contain biological information associated with them are colored in orange, while the ones that were not enriched are in blue. ns > 0.05; 0.001 < * < 0.05; 0.0001 < ** < 0.001; *** < 0.0001

We can see that the coindex managed to identify the largest connected components and also has the most components with biological information. Pearson’s r is the second best in this regard, while Spearman’s is the least effective. Even though the algorithm was run to search for 20 components simultaneously, CoIndex only found 11, Pearson found 14, and Spearman 15. Most of these components comprise a very small number of genes that don’t provide any insight into the co-expressed functions of the enzymes. These small components are also the ones that contain the smallest Z values that resulted in insignificant FDR-corrected p-values.

The images from Figure 7 illustrate the configuration of the subnetworks with enriched genes for each of the three measurements.

**Figure 7:**
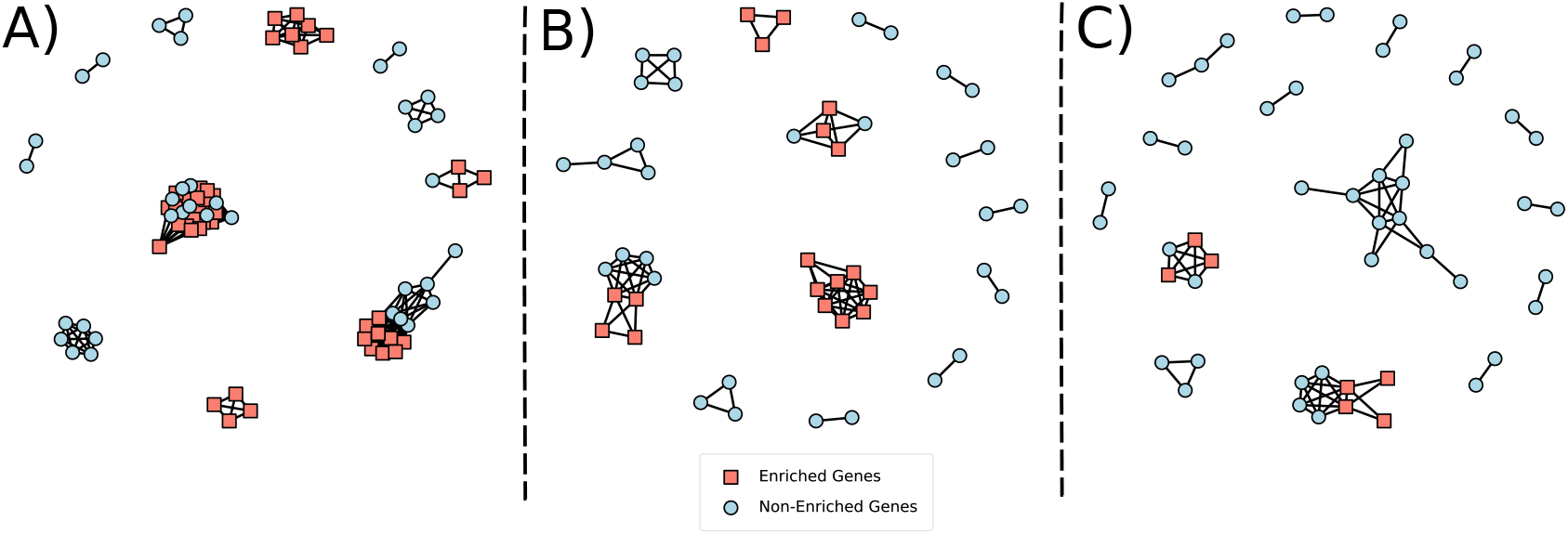
Topology of the enriched components found using all three measurements. A) Components obtained using the coincidence index. B) Components obtained using Pearson’s r. C) Components obtained using Spearman’s r.

Upon examining the genes present in the components of each one of the measurements, we can check that there is some overlap between them (Table 3). The pathway that was present in all three was the cobalamin biosynthesis pathway. Cobalamin is a protein co-factor more commonly known as vitamin B12 Warren et al. (2002). This co-factor is associated with enzymes responsible for the degradation of amino acids and carbon fixation Allen (2012), another pathway that was highlighted by CoIndex and Pearson (but not Spearman).

**Table 3:**
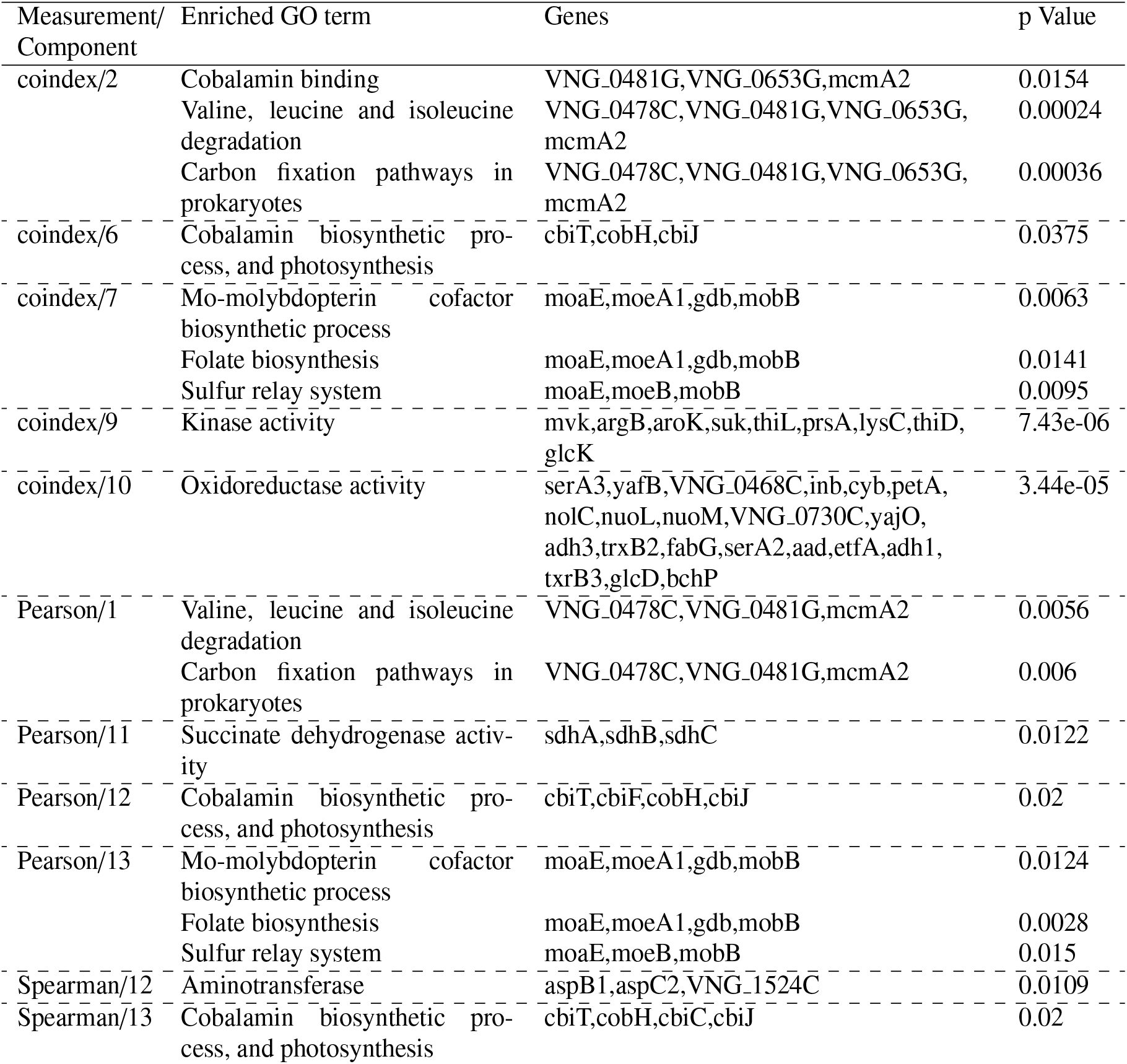
Most relevant enriched biological terms separated by their respective components. The *p* Value is adjusted using FDR.

| Measurement/<br>Component | Enriched GO term | Genes | p Value |
| --- | --- | --- | --- |
| coindex/2 | Cobalamin binding | VNG_0481G,VNG_0653G,mcmA2 | 0.0154 |
|  | Valine, leucine and isoleucine degradation | VNG_0478C,VNG_0481G,VNG_0653G,mcmA2 | 0.00024 |
|  | Carbon fixation pathways in prokaryotes | VNG_0478C,VNG_0481G,VNG_0653G,mcmA2 | 0.00036 |
| coindex/6 | Cobalamin biosynthetic process, and photosynthesis | cbiT,cobH,cbiJ | 0.0375 |
| coindex/7 | Mo-molybdopterin cofactor biosynthetic process | moaE,moeA1,gdb,mobB | 0.0063 |
|  | Folate biosynthesis | moaE,moeA1,gdb,mobB | 0.0141 |
|  | Sulfur relay system | moaE,moeB,mobB | 0.0095 |
| coindex/9 | Kinase activity | mvk,argB,aroK,suk,thiL,prsA,lysC,thiD,glcK | 7.43e-06 |
| coindex/10 | Oxidoreductase activity | serA3,yafB,VNG_0468C,inb,cyb,petA,nolC,nuoL,nuoM,VNG_0730C,yajO,adh3,trxB2,fabG,serA2,aad,etfA,adh1,txrB3,glcD,bchP | 3.44e-05 |
| Pearson/1 | Valine, leucine and isoleucine degradation | VNG_0478C,VNG_0481G,mcmA2 | 0.0056 |
|  | Carbon fixation pathways in prokaryotes | VNG_0478C,VNG_0481G,mcmA2 | 0.006 |
| Pearson/11 | Succinate dehydrogenase activity | sdhA,sdhB,sdhC | 0.0122 |
| Pearson/12 | Cobalamin biosynthetic process, and photosynthesis | cbiT,cbiF,cobH,cbiJ | 0.02 |
| Pearson/13 | Mo-molybdopterin cofactor biosynthetic process | moaE,moeA1,gdb,mobB | 0.0124 |
|  | Folate biosynthesis | moaE,moeA1,gdb,mobB | 0.0028 |
|  | Sulfur relay system | moaE,moeB,mobB | 0.015 |
| Spearman/12 | Aminotransferase | aspB1,aspC2,VNG_1524C | 0.0109 |
| Spearman/13 | Cobalamin biosynthetic process, and photosynthesis | cbiT,cobH,cbiC,cbiJ | 0.02 |

One notable aspect of the coindex measurement is evident in its component 2, where 3 cobalamin binding genes were present (VNG0481G, VNG0653G, mcmA2). This component is analogous to component 1 of the Pearson measurement, but it did not include the cobalamin binding term due to its smaller size, which included only two out of the three cobalamin binding genes. However, both the coindex (component 2) and Pearson’s r (component 1) components had very low Z values. In a situation where this biological function is not known, this might be discarded as statistically insignificant.

The coindex measurement also presents disadvantages. Its two largest and most significant components, 9 and 10, contain genes related to kinase and oxidoreductase activity respectively. These reactions are dependent on metabolites such as phosphates and NADH/NAD+, which are extremely common metabolites that are present in many reactions with different biological functions. This means that the information obtained from these enzymes is too generic, and doesn’t provide a clearer understanding of the possible pathways present in the *H. salinarum* metabolism. The experiments done in the metabolic network with the more common metabolites removed will show how each measurement was affected and how they can reveal more specific biological phenomena.

The second experiment identified around 3 times more components than the first (Figure 8). However, many of them are still very small. Since the FDR correction is directly correlated with the number of instances, many of them didn’t have a high enough Z value to get a significant p-value. This resulted in all of the components identified by the Pearson measurement being considered non-significant (Figure 9). Even though Pearson obtained a similar total Z value as Spearman, each component individually was not able to pass the FDR test. Spearman managed to identify some significant components, but coindex shows the biggest improvement, with 7 large and enriched subnetworks that managed to elucidate more clearly the chemical pathways of the archaeon.

**Figure 8:**
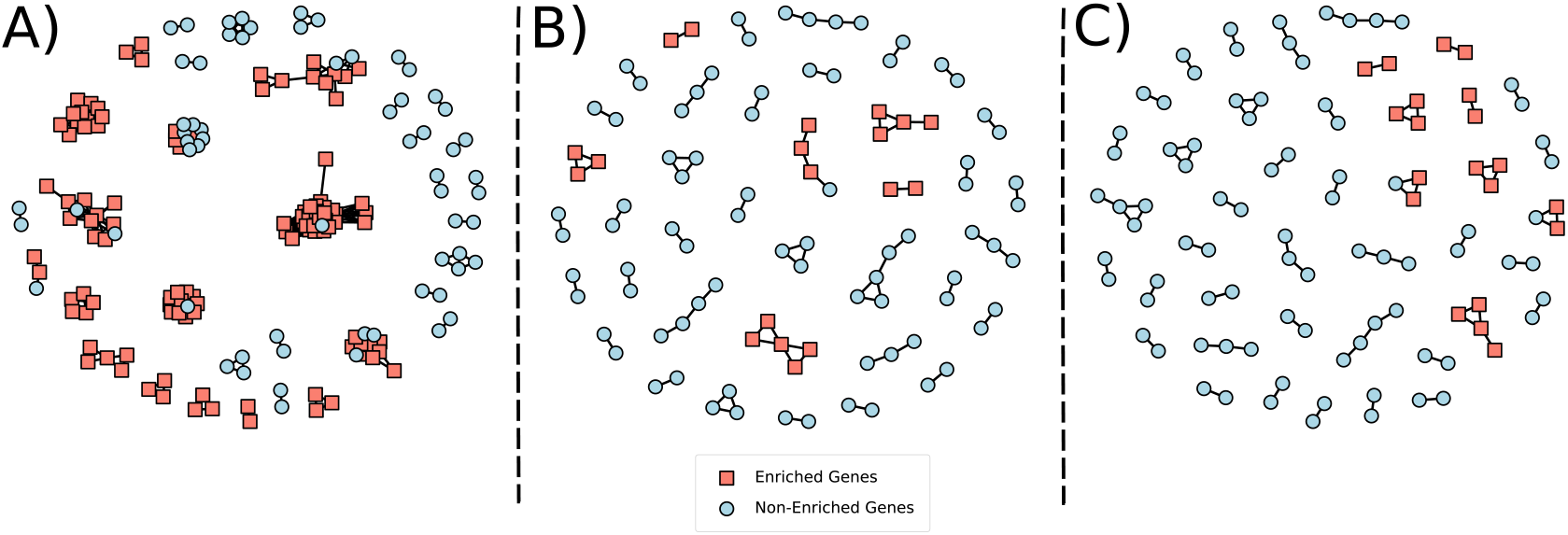
Topology of the enriched components found using all three measurements. This time, using the graph with the top 10 most common metabolites removed. A) Components obtained using the coincidence index. B) Components obtained using Pearson’s r. C) Components obtained using Spearman’s r.

**Figure 9:**
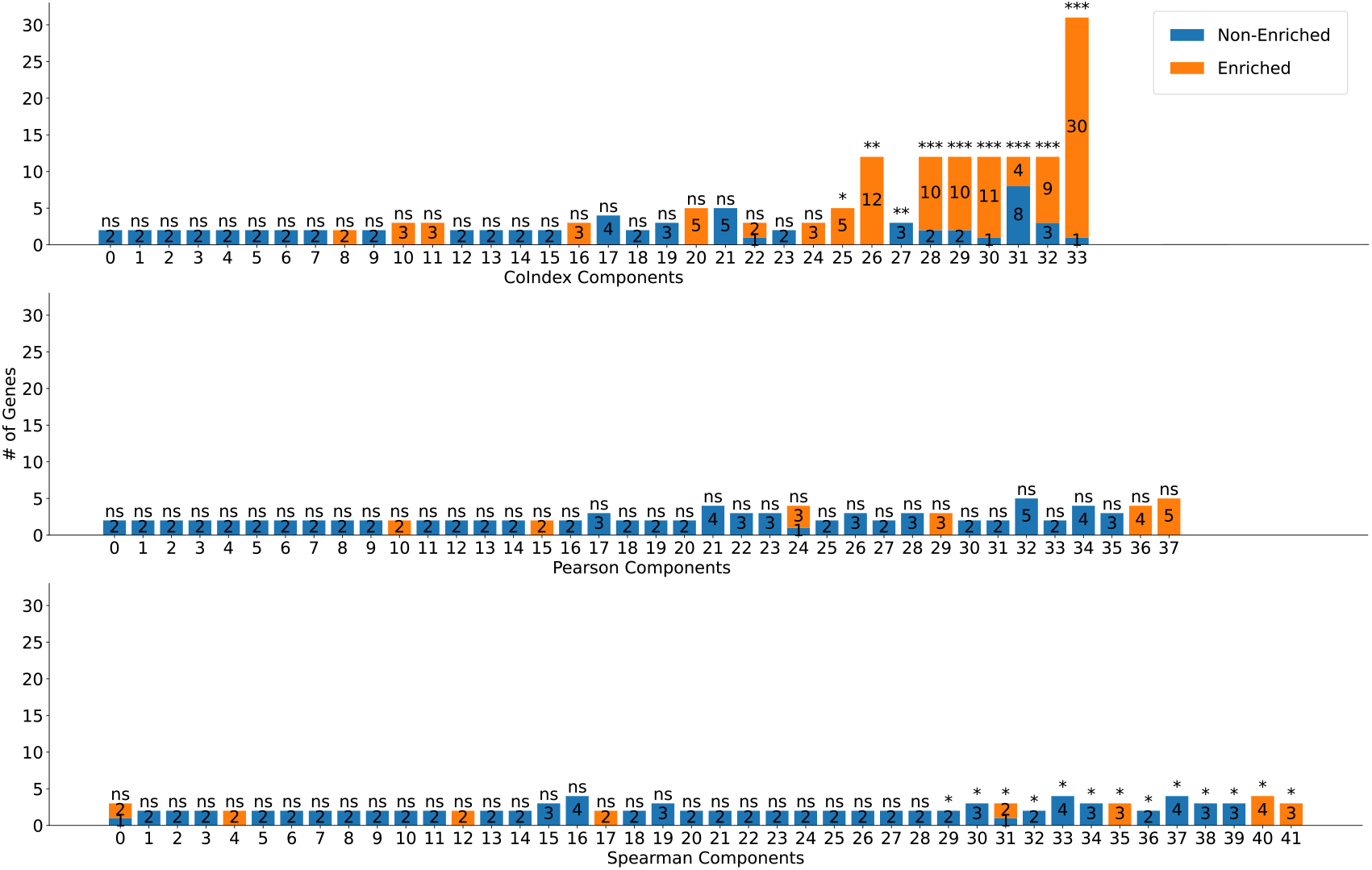
Barplots representing the number of genes contained in each connected component found for the graph without the most common metabolites. It is clear that the coindex got the most enriched components overall, and they also tend to contain more genes than Pearson’s r and Spearman’s r. ns > 0.05; 0.001 < * < 0.05; 0.0001 < ** < 0.001; *** < 0.0001

We hypothesize that coindex was able to achieve the best results because of its stringency rather than despite it. Pearson and Spearman may give higher scores for individual relationships between genes, but this ends up favoring small, highly correlated clusters that do not allow for a reasonable interpretation of the co-expressed pathways. Coindex penalizes noisy data, non-linearity, and variations in scale more harshly and this lowers the score of every relationship significantly, avoiding the creation of ”peaks” with few genes. Instead, this new measurement identifies local maxima that are broader, which include more genes and allows for a more complete picture of the enzymatic reactions.

Upon examining the enriched terms (Table 4), we can now see that the second graph provides a much clearer picture of the metabolic enzymes being co-expressed in the archaeon. We now have the oxidative phosphorylation pathway present. It is responsible for the acquisition of energy metabolites through the electron transport chain (ETC), a combination of transmembrane complexes which generates a proton gradient and forms NADH. The gradient is then used by the ATP synthase to synthesize ATP.Talaue et al. (2016); McKinlay et al. (2020) Moreover, if we take into consideration the experimental conditions that the archaea were exposed to, that is, a continuous anaerobic exposure to light,Baliga et al. (2004) it is safe to assume that anoxygenic photophosphorylation is occurring.McKinlay et al. (2020) In other words, an electron, not derived from hydrolysis with the formation of oxygen, is getting excited by light in a rhodopsin protein and entering the ETC.

**Table 4:**
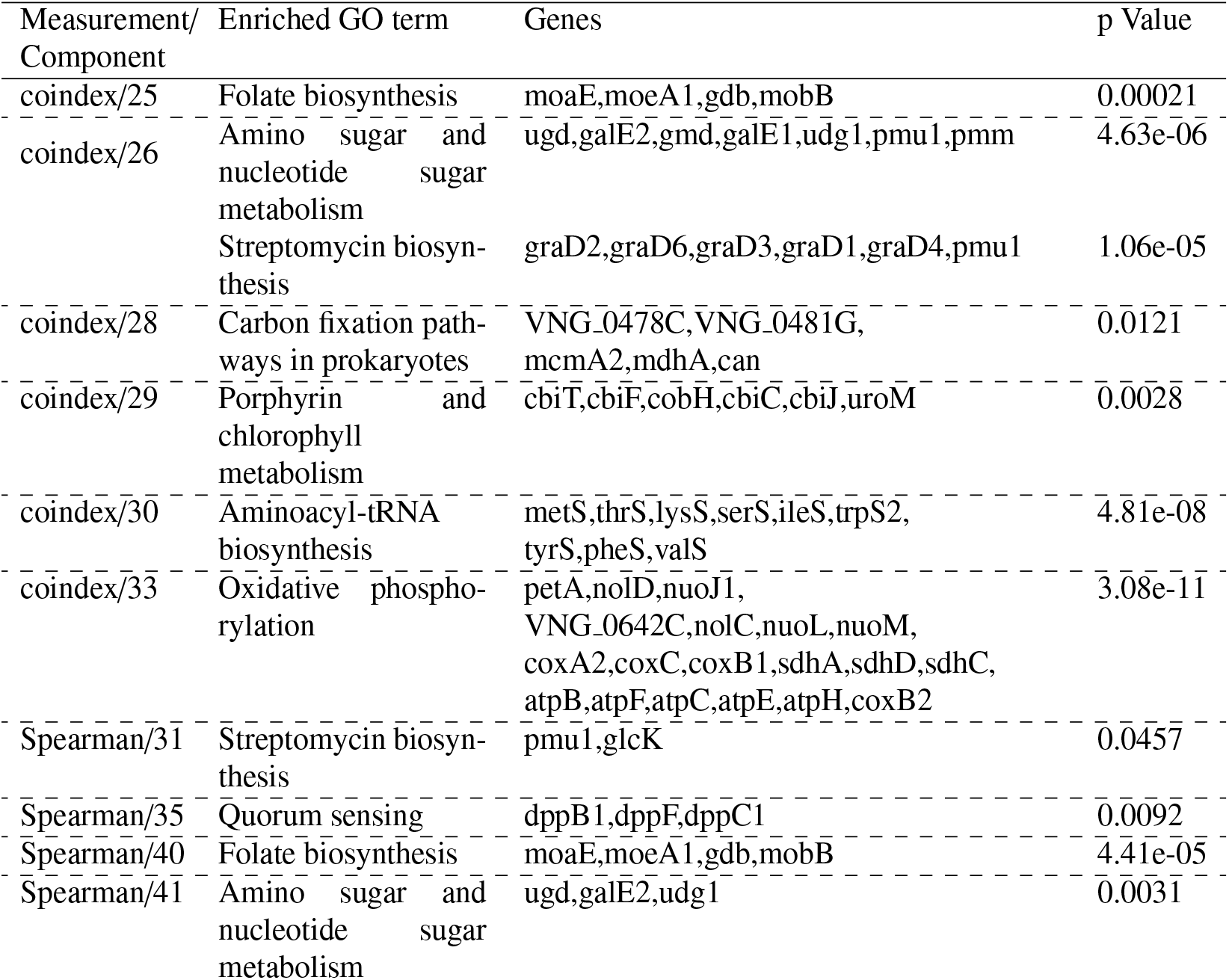
Most relevant enriched biological terms separated by their respective components. Since the second experiment resulted in a lot of components, only the ones that have a significant Z value and enriched KEGG pathways are shown. The *p* Value is adjusted using FDR.

This particular form of oxidative phosphorylation allows *H. salinarum* to extract electrons from molecules with higher potentials such as succinate and acetyl.McKinlay et al. (2020) These molecules can be obtained by the degradation of amino acids, a pathway that is being co-expressed alongside the oxidative phosphorylation one.

Some components that contain terms already present in the first experiment, such as coindex/29, include more biologically correlated genes and yield a more statistically significant result. It is also interesting to note that cobalamin and folate share biological functions and a deficiency in one of them results in a deficiency in the other by a phenomenon called ”methyl folate trap”.Guzzo et al. (2016) This may explain why both of their synthesis pathways are being expressed, because an imbalance in their concentrations can result in a deficiency in both.

Of the four significant components of the Spearman measurement, only one (Spearman/35) contains a unique term. This component contains three genes that code for subunits of a dipeptide transporter with homologs in other prokaryotes known for their quorum-sensing functions.Charlesworth et al. (2017) Spearman/40 contains the same folate biosynthesis genes as coindex/25, but with a more significant result, because it contains fewer genes that are not related to this process. The other two components (Spearman/31 and Spearman/41) contain, separately, many of the same genes present in coindex/26, with the exception of glcK. This demonstrates coindex’s capacity to aggregate smaller components into a more comprehensive and cohesive group.

## 4 Conclusions

As data from transcriptomic experiments become increasingly abundant, more sophisticated methods are necessary to obtain the best possible interpretation of the results. Pearson’s traditional correlation method is widely used in co-expression analysis but, depending on the method utilized, it has significant limitations. In this work, we have shown that, for the discovery of co-expressed genes through enzyme-enzyme interaction graphs based on shared metabolites, the traditional correlation methods, Pearson’s and Spearman’s r, create limited and, sometimes, statistically insignificant subnetworks. The problem appears to worsen with the removal of the most common metabolites, a practice undertaken to avoid biologically unrelated pathways from being mixed up by trivial relationships.

The similarity measurement, coincidence index, proved to be very beneficial for this method. The subnetworks obtained contained more genes, which helped in elucidating more biological pathways and functions and were also more statistically significant. It also showed improved performance with the removal of the common metabolites. In combination with the experimental conditions of the transcriptome experiment, it was possible to conclude that photophosphorylation without oxygen was being induced in the archaeon *Halobacterium salinarum*. It was also possible to identify portions of the pathways related to the synthesis of vitamins, like cobalamin and folate, and amino acid degradation pathways.

Some components included genes that currently do not possess a biological annotation associated with them. Further investigations could be conducted in order to determine whether they are a truly novel addition to the already described pathways or a statistical fluke. Moreover, more recent methods of identifying coexpressed pathways could be tested in order to further evaluate the efficiency of the coincidence index.

